# DrivAER: Identification of driving transcriptional programs in single-cell RNA sequencing data

**DOI:** 10.1101/864165

**Authors:** Lukas M. Simon, Fangfang Yan, Zhongming Zhao

## Abstract

Single cell RNA sequencing (scRNA-seq) unfolds complex transcriptomic data sets into detailed cellular maps. Despite recent success, there is a pressing need for specialized methods tailored towards the functional interpretation of these cellular maps. Here, we present DrivAER, a machine learning approach that scores annotated gene sets based on their relevance to user-specified outcomes such as pseudotemporal ordering or disease status. We demonstrate that DrivAER extracts the key driving pathways and transcription factors that regulate complex biological processes from scRNA-seq data.

## Background

Single cell RNA sequencing (scRNA-seq) experiments dissect biological processes or complex tissues at the cellular and molecular level [1,2]. Due to the high complexity and large number of observations, one critical step in scRNA-seq analysis is dimension reduction [3]. Dimension reduction projects the high dimensional expression matrix into a low dimensional space, also called data manifold or cellular map, which captures the underlying biological processes [4]. A number of methods have been used for manifold learning in scRNA-seq data [5–11].

Biological meaning can be extracted from the data manifold following in-depth analysis. After cells are stratified into separate groups or along a continuum, differential expression analysis is performed. The results can then be examined using tools for biological interpretation [12–14]. However, choosing the best parameters to identify differentially expressed genes across diverse scRNA-seq datasets is still an open challenge [15]. Therefore, there is a need for methods that facilitate biological interpretation without performing differential expression analysis.

Here, we present DrivAER, a method for the identification of Driving transcriptional programs based on AutoEncoder derived Relevance scores. Transcriptional programs (TPs) are sets of genes sharing biological properties [16], which have been annotated extensively [17,18]. DrivAER infers relevance scores for each TP with respect to specified outcomes of interest. These outcomes can represent extrinsic phenotypes, such as disease status, or intrinsic phenotypes derived from the data itself such as pseudotemporal trajectories. Relevance scores allow researchers to rank TPs and help explain the underlying molecular mechanisms.

We evaluated DrivAER by application to two publicly available scRNA-seq datasets and comparison to a competing method called VISION [19]. Our results demonstrate that DrivAER correctly extracts well-known regulators from complex scRNA-seq datasets profiling interferon stimulation and blood development. Our user-friendly tool integrates smoothly downstream of the popular scRNA-seq analysis framework Scanpy [20].

## Results and Discussions

DrivAER is based on one assumption: the data manifold of relevant TPs shares information with the outcome of interest. Irrelevant TPs, on the other hand, will generate data manifolds where the cells fall randomly with respect to the outcome of interest. DrivAER builds upon our Deep Count Autoencoder (DCA) method [10] and iteratively applies DCA to the raw counts of each TP-specific gene set to generate a two-dimensional data manifold in an unsupervised manner (Fig. 1ab). Next, we associate the resulting manifold coordinates with the outcome of interest using random forest models (Fig. 1c). We interpret the random forest accuracy as relevance score, which quantifies the amount of information that is shared between the TP-specific data manifold and the outcome of interest (Fig. 1d).

**Figure 1.**
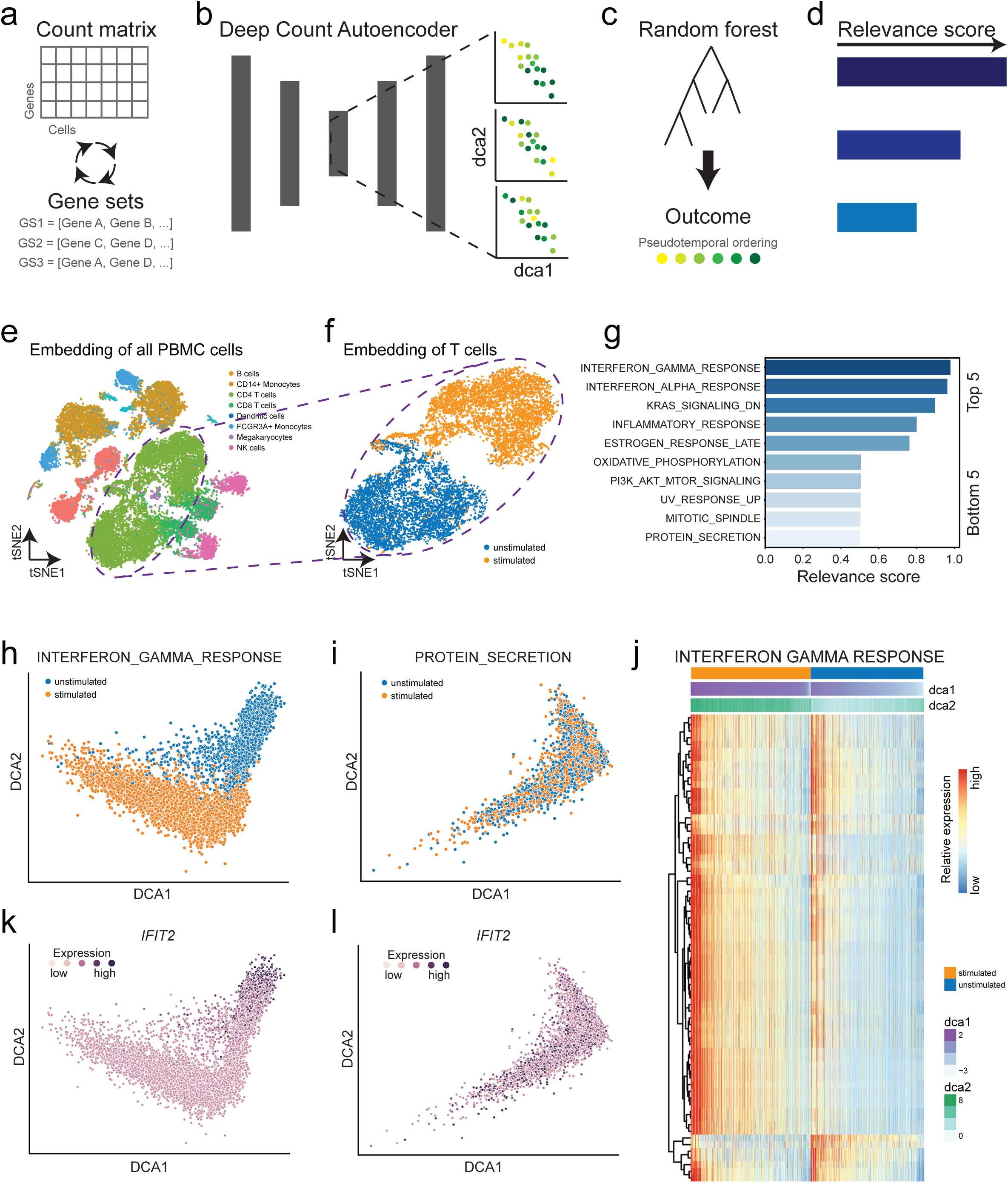
DrivAER correctly identifies interferon response. (a) DrivAER iteratively subjects annotated gene sets to unsupervised dimension reduction via Deep Count Autoencoder (DCA). (b) For each gene set, the generated two-dimensional data manifold coordinates are used as input features in a random forest model to predict the outcome of interest (i.e. pseudotemporal ordering) (c). (d) The random forest prediction accuracy represents the relevance score. (e) t-Distributed Stochastic Neighbor Embedding (tSNE) visualization displays all PBMC cells colored by cell type. (f) Cellular map (tSNE) of T cell subset clusters by stimulation status. (g) Barplot indicates relevance scores of the top and bottom five transcription programs. DCA embedding calculated based on “INTERFERON_GAMMA_RESPONSE” (h) and “PROTEIN_SECRETION” (i) (negative control) gene sets. Cells are colored by stimulation status. (j) Heatmap shows gene expression of “INTERFERON_GAMMA_RESPONSE” target genes and cells in rows and columns, respectively. Columns are ordered first by stimulation status and second by DCA coordinates. Bars on top of heatmap represent stimulation status and DCA coordinates one and two. Red and blue colors correspond to high and low relative expression values. Relative expression of interferon gene IFIT2 is overlaid on top of the DCA embeddings derived from “INTERFERON_GAMMA_RESPONSE” (k) and “PROTEIN_SECRETION” (l) gene sets. Dark colors indicate higher expression.

To demonstrate the ability of DrivAER to perform correct manifold interpretation, we reanalyzed three publicly available scRNA-seq data sets. The first data set by Kang et al [21] described a transcriptional response to interferon stimulation (Fig 1e). described a transcriptional response to interferon stimulation (Fig 1e). As a proof of principle, we asked if DrivAER could recapitulate this biology and extract the interferon signature as the driving transcriptional program defining the T cell data manifold (Fig 1f). We applied DrivAER to the subset of T cells and evaluated 50 hallmark gene sets from MolSigDB [22] with respect to interferon stimulation (Table S1). Indeed, the “INTERFERON_GAMMA_RESPONE” gene set received the highest relevance score (Fig 1g). Visualization of the T cell DCA embedding derived from the “INTERFERON_GAMMA_RESPONE” gene set showed clear separation by condition (Fig 1h), implicating that this gene set is the main driving force separating the stimulated and unstimulated T cells. As a negative control, we show the DCA embedding for one of the lowest scoring gene sets “PROTEIN_SECRETION” (Fig 1i). For this gene set, the cells cluster randomly with respect to the stimulation status. The heatmap in Figure 1j shows the expression levels of T cells for the “INTERFERON_GAMMA_RESPONSE” gene set. It is important to note that the cells (columns) are ordered by stimulation status and that the DCA coordinates are strongly associated with the stimulation status. Most “INTERFERON_GAMMA_RESPONSE” genes are upregulated in stimulated compared to unstimulated cells. Expression of genes in the “PROTEIN_SECRETION” gene set shows a random pattern (Fig S1).

To further manifest the biological meaning of the DCA embedding, we visualized the expression of interferon marker IFIT2 in the “INTERFERON_GAMMA_RESPONSE” (Fig 1k) and “PROTEIN_SECRETION” (Fig 1l) embeddings. Expression levels of IFIT2 increase along the DCA coordinates in the “INTERFERON_GAMMA_RESPONSE” derived embedding. In contrast, IFIT2 expression is distributed randomly in the “PROTEIN_SECRETION” derived embedding.

Next, we tested whether DrivAER is capable of extracting key transcription factors (TF) involved in differentiation trajectories. DrivAER is particularly well suited to infer the relevance of TFs for the following reasons. Firstly, TF mediated regulation is regarded as a combinatorial process which requires the coordination of multiple TFs and co-activators [23]. Secondly, due to the low RNA capture rate in scRNA-seq, generally lowly expressed TFs are not detected reliably [24]. Therefore, the expression levels of the target genes represent a better proxy of TF activity compared to the expression level of the TF itself [25].

To demonstrate the usage of DrivAER, we use a collection of TF-target annotations to infer TF activity and reanalyzed a hematopoietic differentiation dataset by Paul et al [26]. The authors identified and described the main blood development trajectories including differentiation from stem cells towards erythrocytes and monocytes (Fig 2ab). Next, we calculated two independent pseudotemporal trajectories for erythrocyte and monocyte differentiation (Fig 2cd). We then applied DrivAER to identify TFs that are relevant for erythrocyte and monocyte differentiation using the MolSigDB motif gene set [27]. DrivAER identified the GATA TF family as the most relevant in the erythrocyte trajectory (Fig 2e, Table S1). The DCA embedding derived from the “GATA_C” gene set showed strong clustering by pseudotime, demonstrating that GATA target gene expression is highly coordinated along this trajectory (Fig 2f). Indeed, expression levels of GATA targets show strong association with both pseudotime and DCA coordinates (Fig 2g).

**Figure 2.**
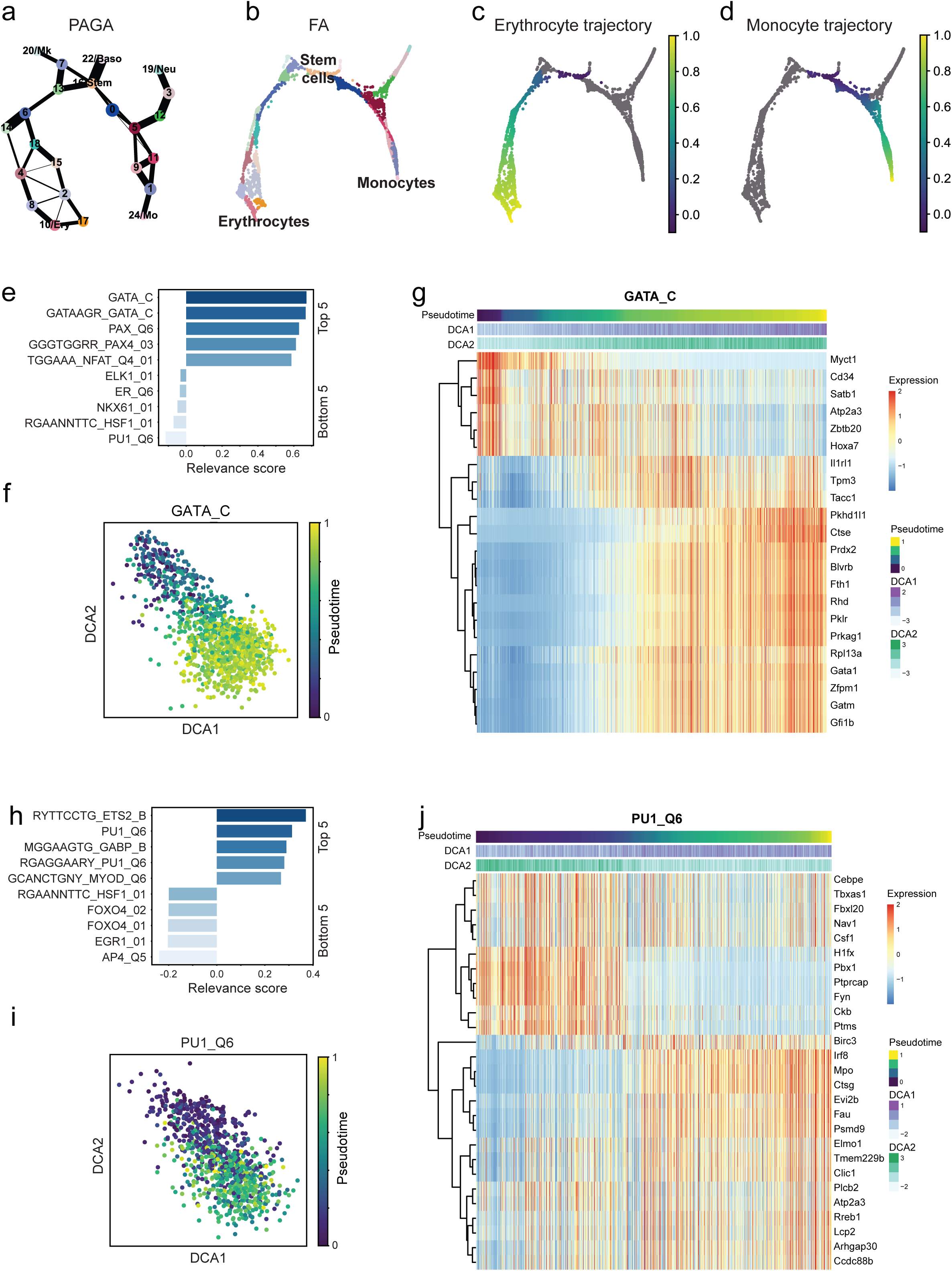
DrivAER unveils key transcription factors in blood development. PAGA (a) and cell-level graph (b) visualization of the Paul et al data set. Cells are colored by Louvain cluster as provided by Scanpy. Two independent trajectories were calculated for erythrocyte (c) and monocyte (d) development. Cells are colored by pseudotime. (e) Barplot displays relevance scores for the 5 most and least relevant transcription factors in the erythrocyte development trajectory. (f) DCA embedding plot was derived from the “GATA_C” gene set and is colored by pseudotime. (g) Heatmap showing gene expression of cells and “GATA_C” target genes for the erythrocyte trajectory in columns and rows, respectively. (h) Barplot displays relevance scores for the 5 most and least relevant transcription factors in the monocyte development trajectory. (i) DCA embedding plot was derived from the “PU1_Q6” gene set and is colored by pseudotime. (j) Heatmap shows scaled gene expression of cells and “PU1_Q6” target genes for the monocyte trajectory in columns and rows, respectively. For both heatmaps, columns are ordered by pseudotime. Bars on top of heatmap indicate pseudotime, DCA coordinates one and two. Red and blue colors reflect high and low expression values.

Of note, targets show both up and down-regulation. A fraction of targets increases in expression along the trajectory while a smaller fraction decreases. When integrating annotation from the TTRUST database [28] with the “GATAAGR_GATA_C” gene set, *Fli1* expression was predicted to be repressed by TF GATA1 and correspondingly we observed a negative correlation along the trajectory between these two genes (Fig S2).

Among the most relevant TFs in the monocyte trajectory was PU1 (Fig 2h, Table S1), which also showed strong association between the DCA embedding (Fig 2j) and target gene expression (Fig 2i) with pseudotime. Both GATA and PU1 are well-known lineage determining regulators in blood development, with GATA and PU1 driving erythrocyte and monocyte differentiation, respectively [29]. However, such conclusions cannot be drawn based on the expression of *Gata1* and *Pu1* itself. Although *Gata1* and *Pu1* showed increased expression along their developmental trajectories, many other TFs exhibited a similar pattern, making it difficult to pinpoint the driving regulator (Fig S3).

Furthermore, to validate our findings we repeated the analysis in a second, independent blood development scRNA-seq data set [30]. In this second data set, DrivAER correctly identified the gene sets “GATA_C” and “PU1_Q6” as the most relevant TFs for erythrocyte and monocyte differentiation, respectively. Taken together, our findings demonstrate that DrivAER robustly explains the molecular mechanisms underlying complex biological processes.

We compared DrivAER to VISION [19] for all three analyses (Fig S4). We observed strong correlation between the relevance scores derived from DivAER and the signature scores derived from VISION for the interferon stimulation experiment, confirming the validity of both methods (Pearson’s correlation, Rho 0.86, p <1e-10). While no significant correlation was observed in the blood development experiments, both methods correctly identified GATA and PU1 related gene sets among the highest scoring TFs for the erythrocyte and monocyte trajectories, respectively. However, VISION scored GATA and PU1 highly in both trajectories, whereas high DrivAER relevance scores were specific to the corresponding trajectories, indicating increased specificity.

## Conclusions

Unlike VISION, DrivAER does not require a pre-defined distinction between the sign of regulation (repression or activation) of genes in a given gene set. The unsupervised nature of the DCA embedding captures any form of non-random, coordinated expression pattern. Therefore, DrivAER captures complex, non-linear expression patterns commonly observed in scRNA-seq data. In principle, the DCA step can be replaced with any other dimension reduction method and future work will focus on potential extensions of DrivAER by integrating different dimension reduction approaches. As illustrated in the blood development example we divided the manifold into independent trajectories for interpretation. However, DrivAER provides the flexibility to be applied to the entire manifold or any subset of it. The user can make this choice and arbitrarily define regions of the manifold, which are expected to be regulated by a TP.

In summary, specialized methods facilitating the functional interpretation of scRNA-seq data are needed to fuel progress in the field. DrivAER is a novel machine learning approach that is effective for manifold interpretation in scRNA-seq data. Our results demonstrate that relevance scores represent a useful measure to extract driving transcriptional regulators from complex scRNA-seq data sets. DrivAER, including usage tutorial, is freely available from Github (https://github.com/lkmklsmn/DrivAER) and we anticipate broad usage by the community.

## Methods

### Transcriptional program annotations

The Molecular Signatures Database (MolSigDB, v7.0) was used to define transcriptional programs [18]. The Hallmark gene set contained 50 gene sets corresponding to specific well-defined biological processes [22]. The C3 transcription factor targets collection contains 610 genes sets in total, where genes share the same *cis*-regulatory motifs from known TF binding sites in the TRANSFAC (v7.4) [31] database around their transcription start sites [27]. The gene sets with motifs not included in the TRANSFAC database were removed. A total of 495 gene sets were utilized in the blood development study. For mouse scRNA-seq datasets, the gene symbols were converted to mouse homologs before running DrivAER.

### DrivAER

Given a collection of gene sets DrivAER uses the Deep Count Autoencoder (DCA) [10] to calculate a two-dimensional data manifold for each gene set. Autoencoders are neural networks that learn an efficient compression of high dimensional data [32]. One important characteristic that distinguishes DCA from other dimension reduction methods is a scRNA-seq specific noise model. The bottleneck layer captures the compression and represents the data manifold. As a default for DrivAER we set the bottleneck dimension to two neurons. DCA takes raw count matrix as input and outputs the data manifold coordinates using the parameter mode = “latent”. To account for differences in library size, size factors derived from the transcriptome-wide expression matrix are calculated and defined in the AnnData object.

The relevance scores are derived using random forest models as implemented in the python module sklearn (v0.21.2) and the two-dimensional data manifold as input features and the variable of interest as outcome. The number of trees was set to 500. The Out-of-Bag accuracy score of the TP-specific random forest model represents the relevance score.

### Interferon stimulation analysis

The scRNA-seq data set of 29,065 PBMCs from lupus patients with and without interferon stimulation were obtained from the GEO database (GSE96583). The t-Distributed Stochastic Neighbor Embedding (tSNE) coordinates in the dataset were used to create the embedding plot, as well as cell type and cell state (stimulated or unstimulated). CD4 T cells were isolated and DBSCAN clustering algorithm [33] was utilized to remove outlier cells (epsilon = 0.1, min_cells = 20). Before applying DrivAER method, lowly expressed genes and cells with less than 3 counts were filtered out.

### Blood development analysis

Expression data for the Paul et al data set was obtained from Scanpy [20] builtin datasets using the “*scanpy.datasets.paul15()”* function. Expression data consists of 2730 hematopoietic stem cells and 3451 genes. The 1000 most highly variable genes were selected for downstream analysis. Louvain clustering, cell type annotation, and diffusion pseudotime calculation was performed following the Scanpy tutorial. Two major developmental trajectories were identified, namely the differentiation of hematopoietic stem cells to erythrocytes and monocytes. Pseudotemporal ordering was independently calculated for these two trajectories using the “*scanpy.tl.dpt”* function. DrivAER was applied to the raw counts and pseudotemporal ordering of each trajectory independently to infer relevant TPs.

Expression data for the Nestorowa data set was obtained from the “Gene and protein expression in adult hematopoiesis” website (http://blood.stemcells.cam.ac.uk/single_cell_atlas). Based on the provided annotation, cells were divided into the erythrocyte and monocyte trajectory. Pseudotemporal ordering was calculated as described above. DrivAER was applied as described above.

### VISION

VISION (version 2.0.0) was downloaded from github (https://github.com/yoseflab/VISION). VISION incorporating a trajectory mode was run on the blood development data following the pipeline of the VISION tutorial. After filtering and normalization, slingshot from the Dynverse package [34] (https://github.com/dynverse/dyno) was used to infer the trajectory and the milestone network. The gene signatures selected by VISION were compared to our results.

## Abbreviations

DCA: deep count autoencoder
MolSigDB: the Molecular Signatures Database
scRNA-seq: single cell RNA sequencing
TF: transcription factor
TP: transcriptional program
tSNE: t-distributed Stochastic Neighbor Embedding

## Declarations

### Ethics approval and consent to participate

Not applicable

### Consent for publication

Not applicable

### Availability of data and materials

DrivAER is available via Github (https://github.com/lkmklsmn/DrivAER).

### Authors’ contributions

LS conceived the idea and designed the project. LS and FY analyzed the data. ZZ supervised the project. LS, FY, ZZ wrote the manuscript. All authors read and approved the final manuscript.

## Acknowledgements

The authors would like to thank the members of the Bioinformatics and Systems Medicine Laboratory at the University of Texas Health Science Center at Houston for stimulating discussion.

