## Supplemental Figures for "DrivAER: Identification of driving transcriptional programs in single-cell RNA sequencing data"

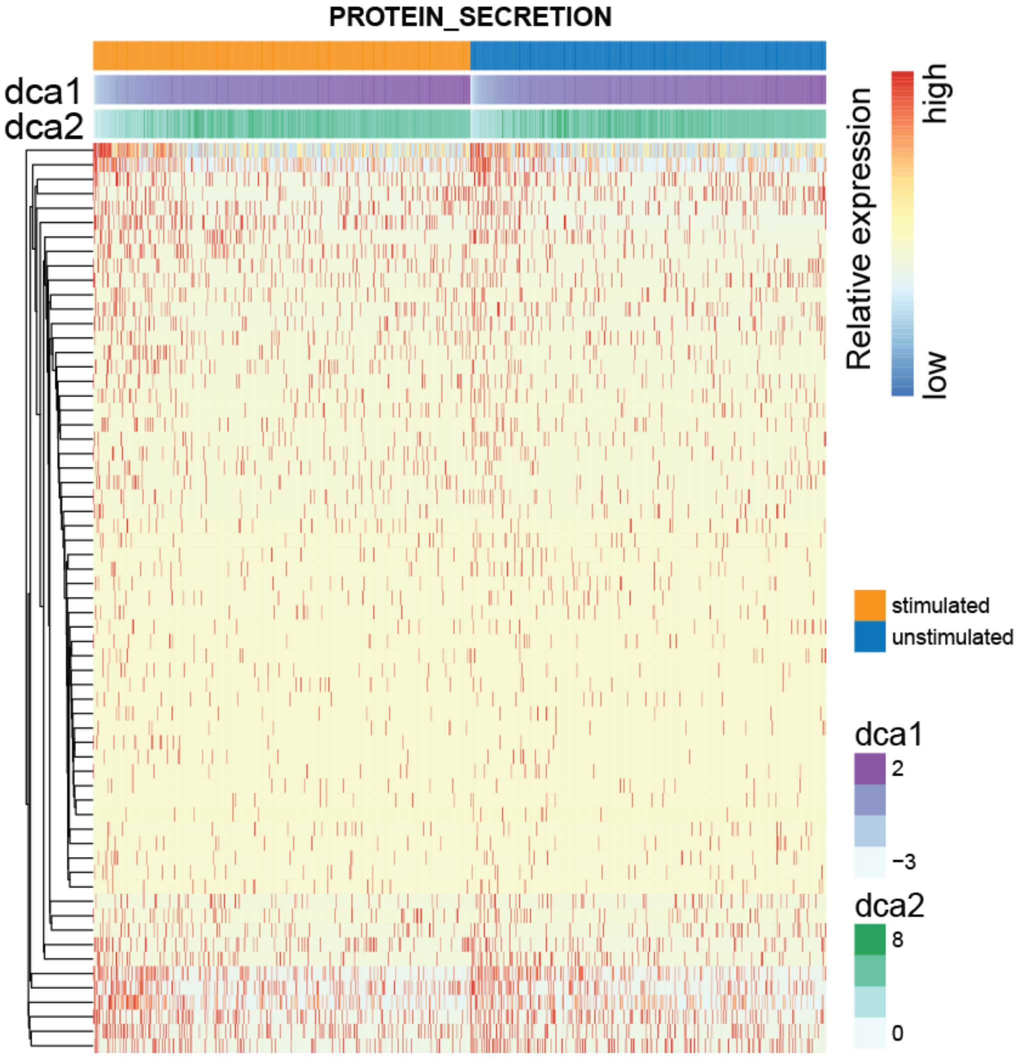


**Figure S1**. Heatmap shows gene expression of PROTEIN_SECRETION gene set and cells in rows and columns, respectively. Columns are ordered first by stimulation status and second by DCA coordinates. Bars on top of heatmap represent stimulation status and DCA coordinates one and two. Red and blue colors correspond to high and low relative expression values.

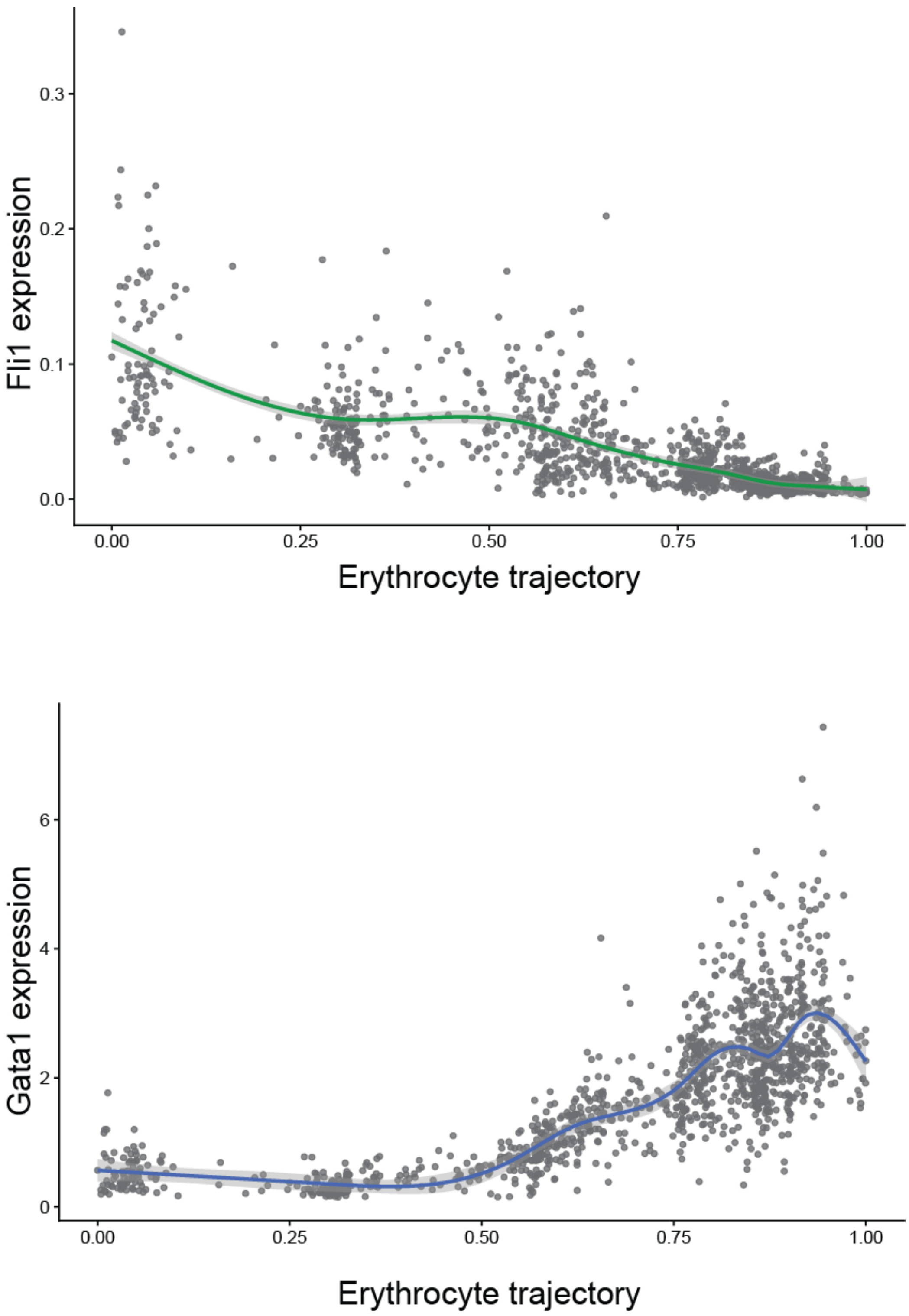


**Figure S2**. Plots show *Fli1* (top) and *Gata1* (bottom) expression along the Erythrocyte trajectory. Grey points indicate cells. The green and blue lines represent smoothed expression estimates.

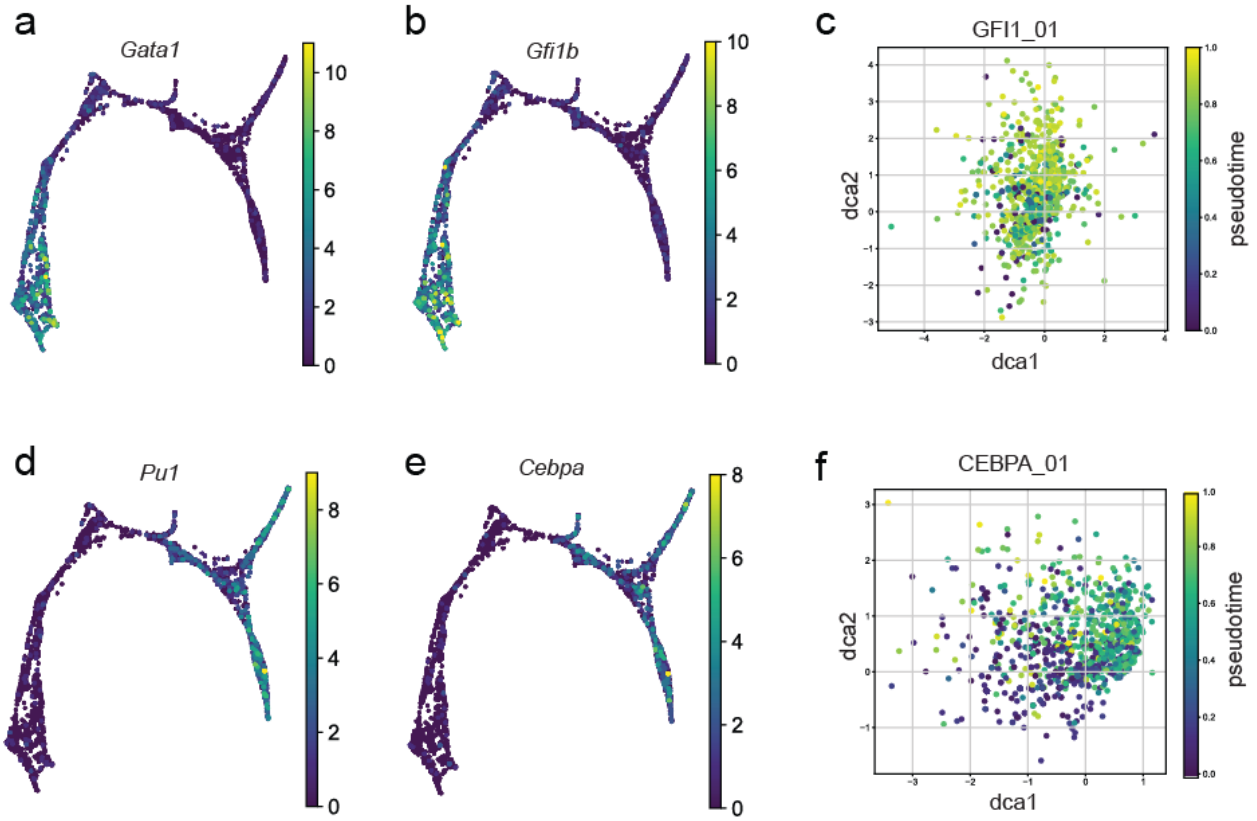


**Figure S3**. Expression of *Gata1* (a) and *Gfi1b* (b) along the erythrocyte trajectory shows similar pattern. However, DCA embedding derived from “GFI1_01” gene set shows poor association with pseudotime. Expression of *Pu1*(a) and *Cebpa*(b) along the monocyte trajectory shows similar pattern. However, DCA embedding derived from “CEBPA_01” gene set shows poor association with pseudotime.

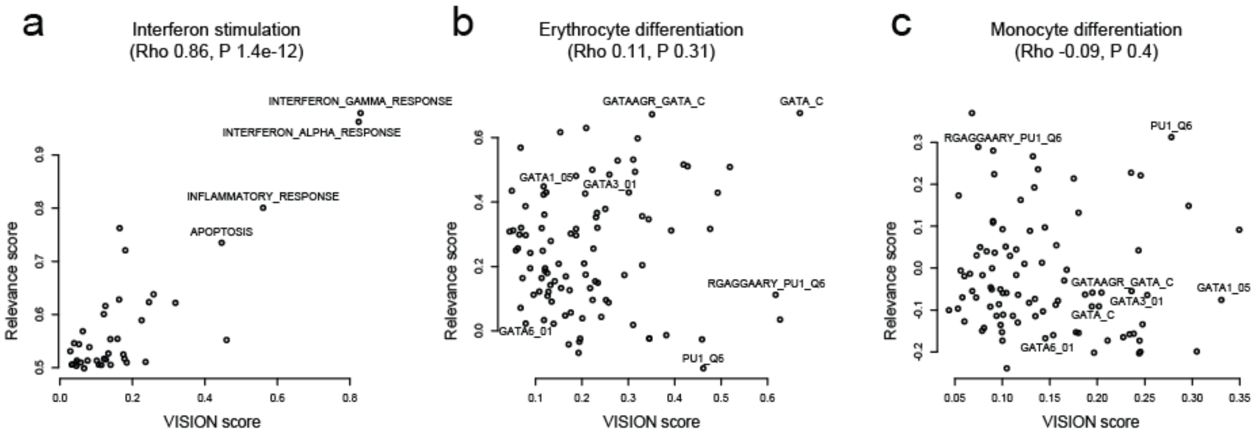


**Figure S4**. Scatter plots depict VISION signature and DrivAER derived relevance score for the Interferon stimulation (a), erythrocyte (b) and monocyte (c) trajectories on X and Y axes, respectively. Points represent gene sets. Exemplary gene sets are highlighted.
